# One-step RNA extraction for RT-qPCR detection of 2019-nCoV

**DOI:** 10.1101/2020.04.02.022384

**Authors:** Monica Sentmanat, Evguenia Kouranova, Xiaoxia Cui

## Abstract

The global outbreak of coronavirus disease 2019 (COVID-19) has placed an unprecedented burden on healthcare systems as the virus spread from the initial 27 reported cases in the city of Wuhan, China to a global pandemic in under three month[1]. Resources essential to monitoring virus transmission have been challenged with a demand for expanded surveillance. The CDC 2019-nCoV Real-Time Diagnostic Panel uses a real-time reverse transcription polymerase chain reaction (RT-PCR) consisting of two TaqMan probe and primer sets specific for the 2019-nCoV N gene, which codes for the nucleocapsid structural protein that encapsulates viral RNA, for the qualitative detection of 2019-nCoV viral RNA in respiratory samples. To isolate RNA from respiratory samples, the CDC lists RNA extraction kits from four manufacturers. In anticipation of a limited supply chain of RNA extraction kits and the need for test scalability, we sought to identify alternative RNA extraction methods. Here we show that direct lysis of respiratory samples can be used in place of RNA extraction kits to run the CDC 2019-nCoV Real-Time Diagnostic assay with the additional benefits of higher throughput, lower cost, faster turnaround and possibly higher sensitivity and improved safety.

## INTRODUCTION

Recent epidemiological models predict a key driver for containing infections within a community is frequent, weekly COVID-19 testing with a rapid reporting turnaround[2,3]. When combined with social distancing, frequent hand-sanitizing, and face mask requirements, these measures have the potential to eradicate transmission over time [4–6]. However, a major bottleneck to expanded COVID-19 testing by RT-PCR is RNA extraction, because this step relies on the availability of consumables, with proprietary compositions, supplied by a limited number of manufacturers.

In the CDC protocol, specimens are typically placed in 3 mL of viral transport media consisting of a buffered salt solution with fetal bovine serum and an antimicrobial cocktail [7]. Viral particles in the samples remain infectious until lysed during RNA extraction. The RNA is then prepared by using a column-based RNA extraction kit. There are four manufacturers with CDC approved RNA extraction kits – Qiagen, Roche, Promega and bioMerieux[8]. While many of these kits are compatible with high-throughput liquid handling platforms, the consumables are expensive and protocols can add an hour or more to the testing schedule[9].

We showed previously that RNA can be extracted by a one-step lysis in QuickExtract DNA Extraction Solution (Lucigen, QE buffer) and directly used in RT-PCR[10]. The QE buffer contains detergents and proteinase K, both of which could inactivate viral particles. Previous work on hepatitis C and Ebola virus has shown that detergent alone is sufficient to reduce infectious titer in the absence of serum (e.g. fetal bovine serum) and is even more effective in combination with proteinase K, which degrades core viral proteins accessible through lipid viral envelop dissolution by detergent[11,12].

Here, we show RNA extraction from nasopharyngeal (NP) and oropharyngeal (OP) specimens by one-step lysis using QE buffer is suitable for the 2019-nCoV RT-PCR assay. We also describe a homebrew one-step lysis buffer that performs comparably well. Importantly, we show one-step lysis buffers maintain sample stability at ambient temperature overnight and support detection of N transcript COVID-19 RNA at 100 copies of input, sensitivity required during early stages of infection and viral reproduction[13–16].

## RESULTS

To test the compatibility of QE extracted samples with the 2019-nCoV RT-PCR assay, we placed self-collected nasopharyngeal swabs directly in 200 ul of QE buffer. Prior to heat extraction, samples were vortexed and divided into two. One was used to create a positive control counterpart with 2019-nCoV plasmid DNA template added at 100 copies/ul. Although 2019-nCoV is an RNA virus, this positive DNA control ensures the assay is working while maximizing safety and the need for Biosafety Level 2 (BSL2) precautions, as is required for handling 2019-nCoV RNA material. An RNA extraction control using HCT-116 human colorectal cancer cells at 10 cells/ul was included. Samples were then heated at 65°C for 15 min followed by 98°C for 2 min to inactivate proteinase K and then directly used for the test using a single probe and primer set (N1) as well as the set for RNaseP.

All positive control counterpart samples had a Ct value for N1 between 29-32, while all experimental samples had no detectable amplification from the N1 primer/probe set (Figure 1). We detected RNaseP reference signals in both the OP and human cell line extraction controls, which confirms that the extractions were successful. Although an equal volume of lysis extract was used to generate the positive control counterpart of OP experimental samples, RNaseP reference signals were 2 cycles greater for positive controls, suggesting that the swab containing portion retained more cells. But it’s important to point out that the raw Ct values do not change the interpretation of results for what is intended to be a qualitative yes/no assay – that the positive and extraction controls performed within the range (<40) to determine the presence or absence of 2019-nCoV N gene in experimental samples[8].

**Figure 1.**
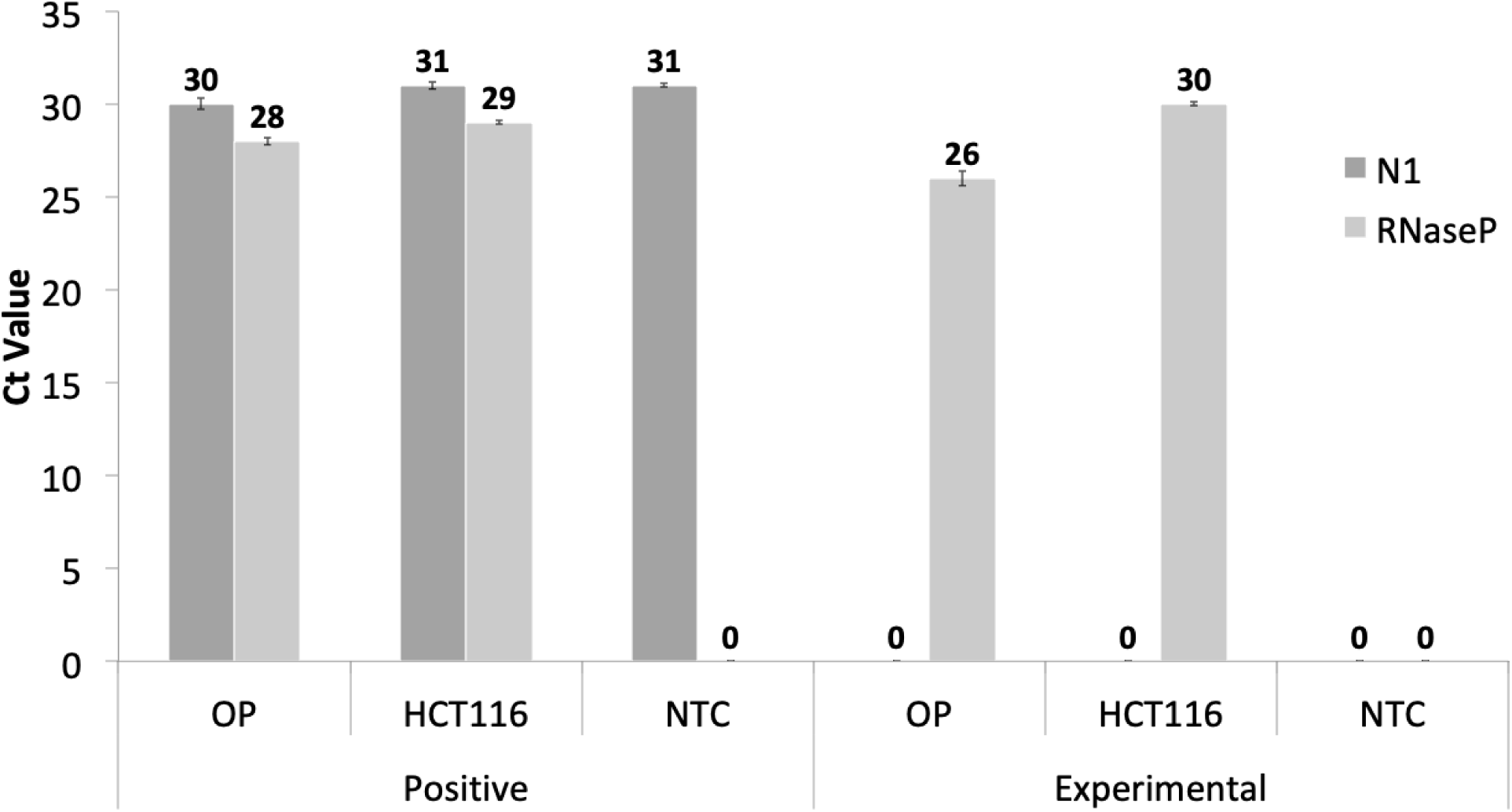
An OP specimen was lysed in QE buffer alongside HCT116 cells as controls and amplified by using one-step RT-PCR. Ct values for OP specimen, HCT-116 extraction control, and no template control containing no cells as input (NTC) shown. A Ct value of 0 indicates no signal was detected. Each sample has a positive control with 100 copies/ul plasmid DNA template spiked-in to confirm N1 probe and primer set performance. Error bars are the standard deviation for three technical replicates.

To determine if NP and OP specimens remain stable in QE buffer until they can be transported from the site of collection to the lab, we stored the samples with positive plasmid DNA template control at 100 copies/ul at room temperature, 4°C, or −20°C for 24 hours. Samples were heat extracted after 24 hours and used for the RT-PCR test with probe sets N1 and N2. The sample without positive control added was processed for extraction immediately after the collection. All positive control samples had a Ct value between 33-34 for N1 and N2 across conditions. The Ct values for reference RNaseP were within the range of 26-29 when the collection swab was present during heat extraction and a Ct range of 29-31 when the swab was not present. This is consistent with epithelial cells being caught in the polyester fibers during collection. The spiked plasmid DNA does not seem to be retained on the swab. The Ct values of samples stored at different temperatures are not significantly different from those samples processed without storage (Table 1).

**Table 1.**
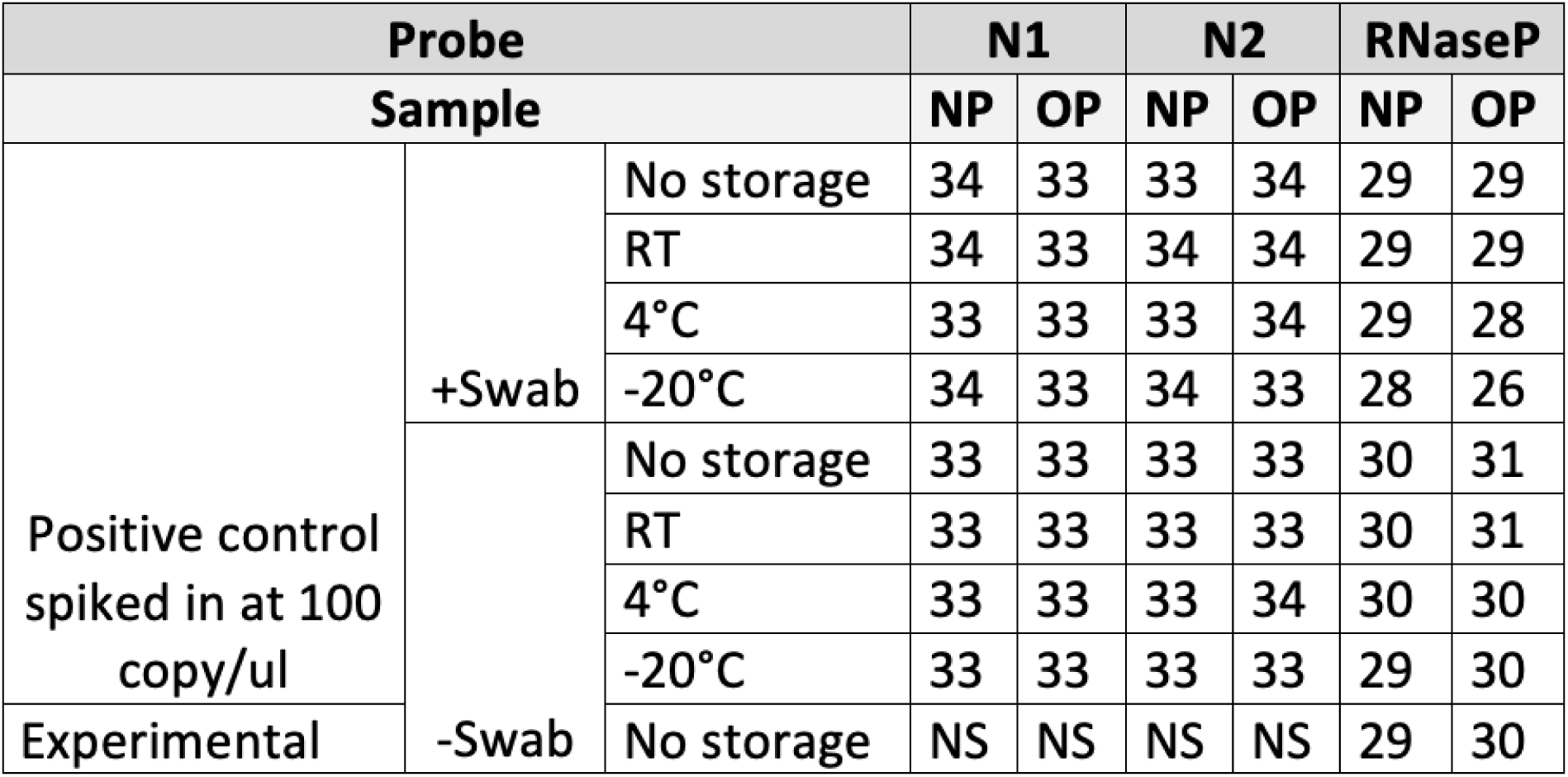
Comparison of QE lysis performance across specimen storage temperatures after 24 hours. Ct values for oropharyngeal (OP) and nasopharyngeal (NP) specimens that were immediately heat extracted (no storage) or kept at room temperature (RT), 4°C or −20°C for 24 hours. The presence of the collection swab during heat extraction is also shown (+/− swab). The positive control is plasmid DNA template spiked at 100 copies/ul into specimen sample. A value of NS indicates no signal was detected.

We sought to compare yields from direct input of QE buffer-lysed sample and column purified RNA. An OP specimen was placed in 200 ul of QE buffer and the collection swab removed. We removed 20 ul for our experimental input before adding 100 copies/ul of positive control DNA template to the remaining buffer. We took 120 ul for column purification using the Qiagen RNeasy Mini Kit with a final elution volume of 30 ul. The remaining 60 ul of QE sample positive control counterpart and 20 ul of experimental QE sample were heat extracted. Each RT-PCR reaction had 2 ul of input assayed. We found QE processed and column purified material had comparable Ct values despite 4-fold more material processed for column purification (Figure 2).

**Figure 2.**
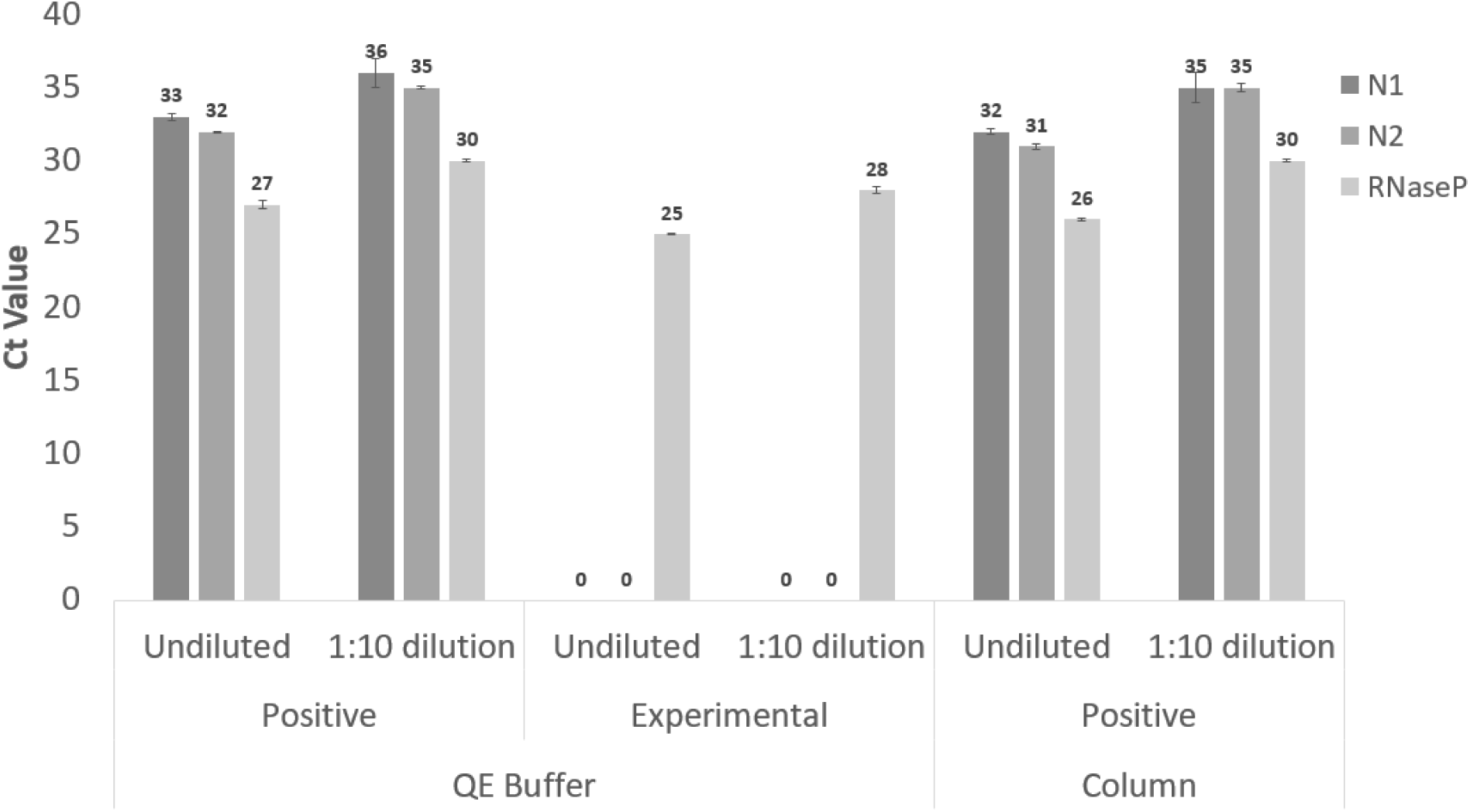
Side-by-side comparison of OP specimen from direct lysis in QE buffer vs. Qiagen RNeasy column purification. Ct values for probe sets N1, N2 and RNaseP shown using undiluted and 1:10 dilutions of positive control counterparts at 100 copies/ul of plasmid DNA template and experimental OP specimen. A Ct value of 0 indicates no signal was detected.

To decrease cost and circumvent supplier bottlenecks, we compared the performance of a home formulated extraction buffer (GEiC buffer) composed of 0.2% Triton and proteinase K with QE buffer using samples spiked with *in vitro* synthesized (IVT) COVID19 N transcript. Here, tongue scrapes were self-collected using sterile polyester flock swabs and placed in 80 ul of either GEiC or QE buffer. Samples were briefly vortexed and four 20 ul aliquots were used to spike-in dilutions of N transcript prior to heat extraction. In each sample, the spiked N transcript copies per RT-PCR reaction were: 1×10^9 (1), 1×10^8 (2), 1×10^7 (3), and 1×10^6 (4). The Ct values for GEiC and QE buffer extracted samples were comparable across dilutions and probe sets, supporting that the GEiC buffer works as well as the QE buffer in maintaining RNA integrity during the extraction process (Figure 4). The IVT N transcript RNA was detectable at a Ct of 35 for 100 copies after heat extraction in GEiC buffer, suggesting RNA stability even at lower copy numbers.

## DISCUSSION

We presented here an improvement to the standard 2019-nCoV RT-PCR test. We replaced the column-based RNA extraction with a one-step lysis. Our indicators for adequate assay performance are consistent with CDC assay interpretation guidelines, the detection of a Ct level of <40 for control 2019-nCoV positive control counterparts and for RNaseP reference signal for positive controls and experimental samples[8]. The direct lysis takes less than 20 min to process samples ready for RT-PCR and can be easily scaled up to a 96-well format and obtain higher throughput. The cost of the lysis buffer is much lower than that of a column purification kit. The lysis only requires a regular PCR machine and does not need a centrifuge or a manifold, as column purification requires.

We demonstrated that both the commercially available QE buffer and our homebrew GEiC buffer work equally well. When necessary, the homebrew buffer can be easily produced in large quantities with minimal cost. This way, it eliminates the potential shortage of RNA extraction kits.

We also demonstrated that the samples are stable in the QE buffer for at least 24 hours even when they are stored at ambient temperatures, allowing time flexibility between sample collection and processing. Detergents and proteinase K in the buffer inactivate the viral particles in positive samples after lysis, making the samples safer to in the testing lab.

In the parallel comparison of QE buffer lysed samples versus column purified samples, the Ct values for positive controls and RNaseP were within 1 cycle of each or less, even though the samples were concentrated 4x (from 120 ul to 30 ul elute). It indicates that there is significant loss of material by column purification and the benefit of concentrating the samples is evened out. Additionally, our method starts with only 200 ul of lysis buffer, and 2 ul of direct lysis is 1/100 of the total sample. We lowered the volume of lysis buffer for the routine test in the lab to 100 ul. With the standard method, the sample transport buffer is 3 mL and only 120 ul to 140 ul, depending on the extraction kit, is taken for purification, essentially a 1:15 dilution that can translate into up to a difference of 4 Ct’s.

This work used DNA and RNA template as a positive control for the assay. Specimens such as NP and OP samples will invariably introduce RNases into the buffer, which during lysis, could degrade viral RNA released after capsid denaturation. The RNA transcript data shows that when free N transcript RNA is added directly into the specimen samples and heat extracted, RNA stability is relatively consistent across extraction buffers (Figure 3), even though we observed some loss of N transcript RNA during extraction. Naked IVT transcripts are likely to be degraded before viral RNA protected by capsid. Our data shows that the assay can detect as few as 100 copies of N IVT RNA transcript after one-step lysis. However, our method has not been tested with known COVID19 positive samples, so sensitivity for such clinical samples remains to be determined.

**Figure 3.**
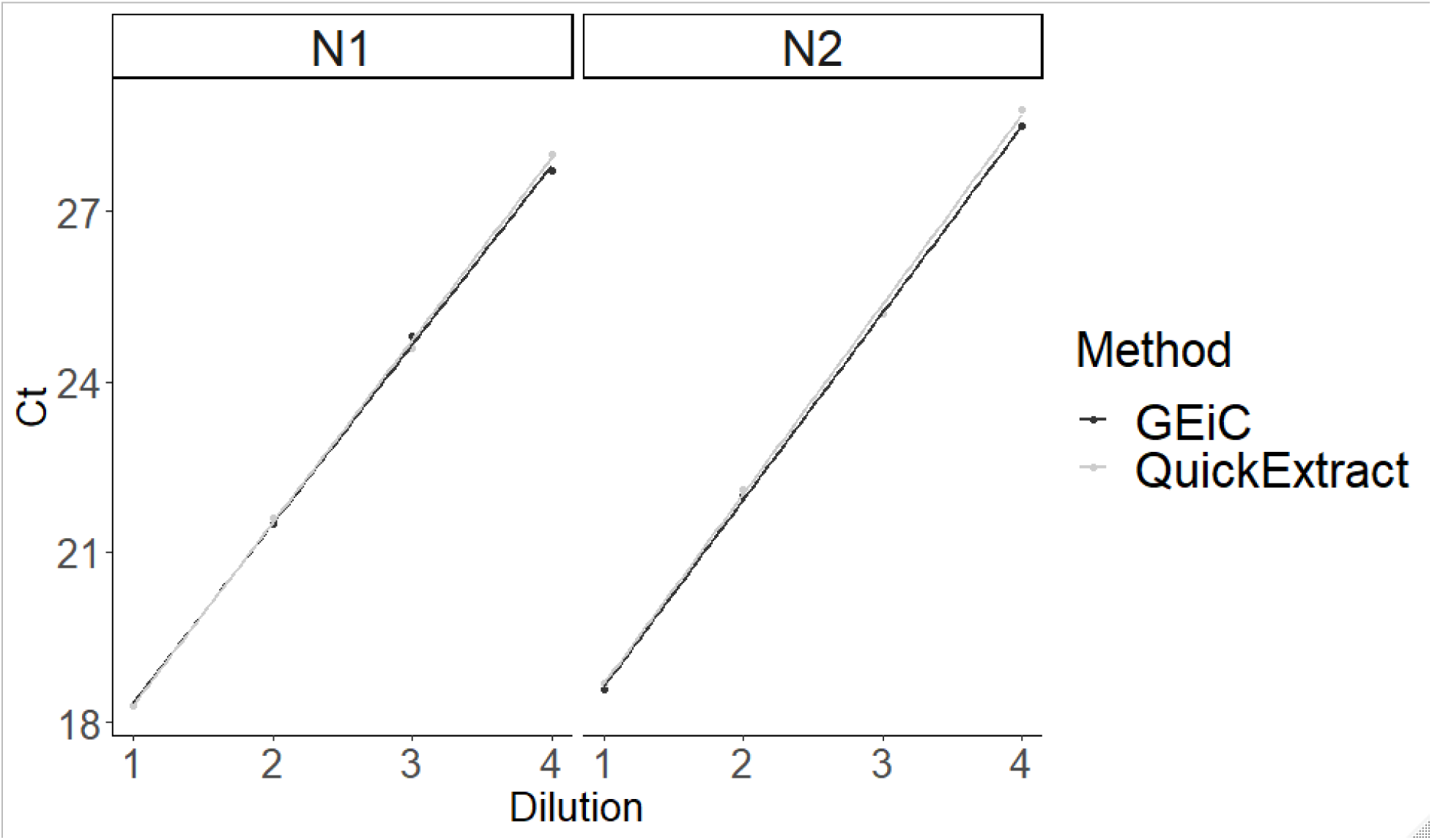
Comparison of GEiC and QE lysis buffer performance. Ct values for tongue swab specimens spiked with 10^10 (1), 10^9 (2), 10^8 (3), or 10^7 (4) copies of N RNA transcript prior to heat extraction using GEiC buffer (dark grey) or QuickExtract (light grey).

**Figure 4.**
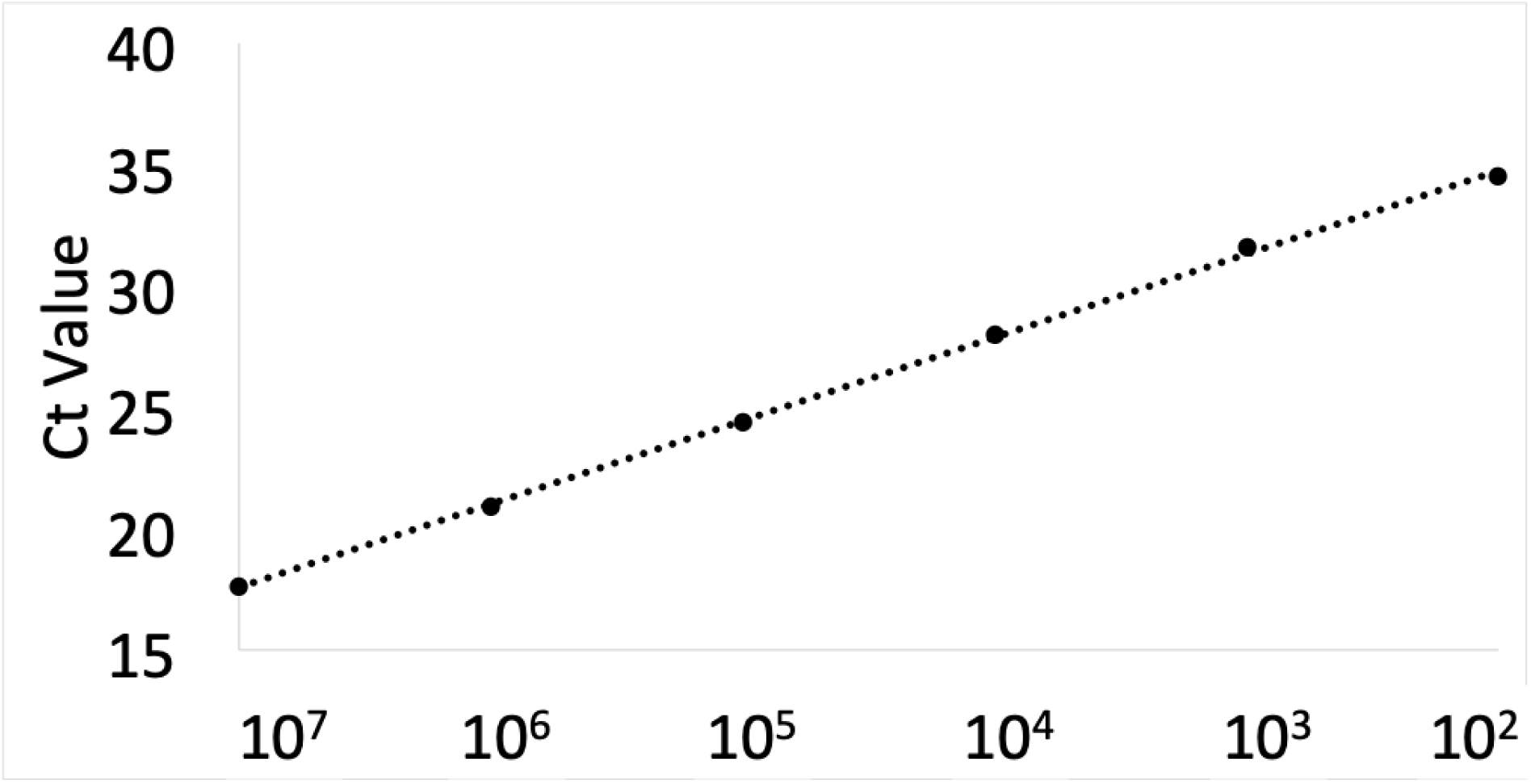
2019-nCoV RT-PCR assay performed on a dilution series of N IVT RNA transcript after heat extraction in homebrew one-step GEiC lysis buffer. The X-axis indicate the total number of copies present in each reaction.

Given the increasing need for more tests to be done quicker, our data presents a feasible option for large scale testing sites. Bypassing the need of column purification and lowering the volume of sample collection buffer together simplify high-throughput process development, shortens turnaround, reduces cost and may improve sensitivity.

We set up a weekly testing schedule in the lab when we returned to work after the shutdown. Everyone may collect, extract and submit their own samples willingly one or more times throughout the week. Extraction inactivates potentially positive samples. Tests are then done by one designated scientist for consistency on all samples each Monday. Even though testing is not a direct means of protection, such as wearing masks, washing hands frequently and keeping physical distance, it has been a nice benefit we enjoy that provides a sense of much needed comfort during this unprecedented time.

## METHODS

### Specimen Collection

Collection of a nasopharyngeal specimen (NP) is the recommended method for testing patients presenting COVID-19 symptoms using the CDC diagnostic RT-PCR test, with oropharyngeal (OP) being an acceptable alternative according to CDC guidelines. To determine if direct lysis of comparable specimen samples could be used for the RT-PCR test, we self-collected NP and OP samples using sterile polyester flock swabs (PurFlock Ultra flocked collection swabs by Puritan Diagnostics LLC). The swabs were each placed in 1.5 mL Eppendorf tube containing 200 ul of QuickExtract DNA Extraction Solution (QE buffer, Lucigen LLC, Madison, WI). Each tube was vortexed and stored until extraction.

Extraction control HCT-116 cells were counted using a hemocytometer and pipetted directly to the QE buffer, and no swab was used. A positive control was made for each sample by transferring 100 ul of sample to a new 1.5 mL Eppendorf tube and adding 10000 copies of COVID19 plasmid DNA template control (IDT Cat. No 10006625), prior to extraction incubation. The final concentration of the positive control is 100 copies/ul. No cell input negative control samples were also used for Figure 1 and Table 2. In Figure 1, the no cell input negative control also had an RNA template positive control counterpart.

### RNA Extraction

Samples were incubated in either GEiC (10 mM Tris, pH 8, 2 mM EDTA, 0.2% Triton X-100, 200 ug/ml proteinase K) or QE buffer at 65°C for 15 minutes, followed by 98°C for 2 min. Column purification for DNA spiked samples was performed using Qiagen RNeasy Mini Kit (Cat. No. 74104) from 120 ul of each sample, using an elution volume of 30 ul.

RNA extraction of personal samples is done by each individual themselves so that all potential positive samples are inactivated before submission, to avoid unnecessary exposure for the designated scientist who performs weekly testing on all samples.

### In vitro RNA synthesis

Template for COVID19 N transcript was amplified from a plasmid (IDT Cat. No. 10006625) using primers nCov-N-T7-F 5’ aaaaTAATACGACTCACTATAGGatgtctgataatggacccca 3’ and nCov-N-R 5’ ttaggcctgagttgagtcag 3’. PCR template was purified with PureLink PCR purification kit (ThermoFisher, Cat. No. K310002).

HiScribe^™^ T7 Quick High Yield RNA Synthesis Kit (NEB, Cat. No. E2050S) was used for in vitro transcription of COVID19 N RNA according to manufacturer’s instructions.

### RT-PCR

Probe and primer sets were obtained from the research COVID19 RT-PCR kits 2019-nCoV RUO (IDT Cat. No. 10006713). A one-step reaction mix was prepared using Reliance One-Step Multiplex Supermix (BioRad Cat. No. 12010220) with 2 ul of sample, 1.5 ul of the probe/prime set mix for each reaction, 5 ul of 4x reaction mix, and 11.5 ul of molecular biology grade water to a final volume of 20 ul. The RT-PCR was run on a QuantStudio^™^ 3 real-time PCR machine (ThermoFisher).

### RECOMMENDED PROTOCOL

#### Pre-aliquoted collection tubes

one microtube with 100 ul of GEiC buffer and two PCR tubes marked A and B. PCR tube A is empty, and PCR tube B has 1 ul of what concentration of IVT of N

#### Sample collection and extraction

1. Each person takes their own NP or OP sample and break the handle, leaving the swab in the microtube.
2. Vortex the microtube and pipet 20 ul each to PCR tubes A and B. Mark both tubes with their own name.
3. Run QE PCR protocol: 65°C, 15 min, 98°C, 2 min on PCR tubes A and B and afterwards place extracted samples in −20C.
4. Dispose the microtube in biohazard waste.

#### qRT-PCR by one designated scientist

1. Take 2 ul each from “+” and “-” samples and assemble the one-step RT-PCR reactions, containing 1.5 ul of each primer/probe set, 5 ul of 4x reaction mix and 11.5 ul of nuclease-free water.
2. Run the reactions on a QuantStudio^™^ 3 real-time PCR machine (ThermoFisher) using the following program: 50°C for 10 min 95°C, 10 min 40 cycles of 95°C, 3 sec and 55°C, 30 sec

#### Data analysis

Data analysis was performed using QuantStudio^®^ Design and Analysis Desktop Software. Example data provided in Supplemental Table 1.

